# Stereotactic electroencephalography in humans reveals multisensory signal in early visual and auditory cortices

**DOI:** 10.1101/549733

**Authors:** Stefania Ferraro, Markus J. Van Ackeren, Roberto Mai, Laura Tassi, Francesco Cardinale, Anna Nigri, Maria Grazia Bruzzone, Ludovico D’Incerti, Thomas Hartmann, Nathan Weisz, Olivier Collignon

## Abstract

Unequivocally demonstrating the presence of multisensory signals at the earliest stages of cortical processing remains challenging in humans. In our study, we relied on the unique spatio-temporal resolution provided by intracranial stereotactic electroencephalographic (SEEG) recordings in patients with drug-resistant epilepsy to characterize the signal extracted from early visual (calcarine and pericalcarine) and auditory (Heschl’s gyrus and planum temporale) regions during a simple audio-visual oddball task. We provide evidences that both cross-modal responses (visual responses in auditory cortex or the reverse) and multisensory processing (alteration of the unimodal responses during bimodal stimulation) can be observed in intracranial event-related potentials (iERPs) and in power modulations of oscillatory activity at different temporal scales within the first 150 ms after stimulus onset. The temporal profiles of the iERPs are compatible with the hypothesis that MSI occurs by means of direct pathways linking early visual and auditory regions. Our data indicate, moreover, that MSI mainly relies on modulations of the low-frequency bands (foremost the theta band in the auditory cortex and the alpha band in the visual cortex), suggesting the involvement of feedback pathways between the two sensory regions. Remarkably, we also observed high-gamma power modulations by sounds in the early visual cortex, thus suggesting the presence of neuronal populations involved in auditory processing in the calcarine and pericalcarine region in humans.

## 1. Introduction

The ability to rapidly and seamlessly integrate information encoded in separate sensory systems is paramount to our ability to adaptively interact with the environment. The brain must, therefore, be endowed with computational mechanisms allowing information originating from multiple sensory systems to be unified into a coherent internal representation of our surroundings (Meredith et al., 1987). The classical view of neocortical organization posits that primary sensory regions process unisensory inputs that are then passed on to association cortices where information from the different senses converge and are integrated. More recently, a plethora of functional imaging (Calvert et al., 2001, Macaluso et al., 2000, Foxe et al., 2002, Martuzzi et al., 2006) and electrophysiological studies in animals and humans (Giard and Peronnet, 1999, Molholm et al., 2002, Raij et al., 2010, Kayser et al., 2008, Lakatos et al., 2007, Brosch et al., 2005, Ghazanfar et al., 2005) provided evidence for multisensory interactions already at the level of early sensory cortices. These studies unhinged the idea that primary sensory areas are exclusively sensitive to sensory input from one modality only, and led to the hypothesis that multisensory integration (MSI) is present in almost all the neocortex, including the earliest cortical stages of sensory processing (Ghazanfar and Schroeder, 2006, Murray et al., 2016).

Even though this new conceptualization has made a pervasive breakthrough in the literature, the debate about the presence of MSI in early sensory regions (Kayser, 2010), as well as the potential underlying mechanisms allowing its implementation, is still far from being resolved (Schroeder and Lakatos, 2009, Mercier et al., 2013, Kayser et al., 2009). Recent findings have just begun to reveal the complexity and the heterogeneity of early MSI (Iurilli et al., 2012, Lakatos et al., 2007, Mercier et al., 2013, 2015). Moreover, recent studies have failed to observe MSI in early sensory regions, further challenging this proposal (Lemus et al., 2010, Quinn et al., 2014).

One important difficulty for the characterization of early multisensory response in regions classically considered as purely unimodal links to the possibility to record brain activity with high spatial and temporal resolution simultaneously (Driver and Noesselt, 2008). For instance, the relative lack of spatial precision of electroencephalographic (EEG) and magnetoencephalographic (MEG) recordings, even when combined with source reconstruction, makes it difficult to convincingly determine if the multisensory responses arise from primary sensory regions or from secondary sensory or higher-level associative regions. Moreover, these techniques are affected by volume conduction problems, making it difficult to assert that the integrative process truly occurs in the early sensory region, since proximate associative regions also engage in MSI and separating nearby source of activity is far from trivial. This also limits the simultaneous exploration of MSI in the auditory and visual cortices (Besle et al., 2004). In contrast, the relative lack of temporal precision of functional magnetic resonance imaging (fMRI) tampers the demonstration of early MSI since the activity can well be explained by feedback connection from heteromodal regions (Goebel and van Atteveldt, 2009) that could lead to reentrant modulation of unisensory responses due to attentional modulation, for instance (Talsma et al., 2010).

In our study, we thwarted these problems by relying on the unique spatiotemporal resolution provided by recording brain activity using intracranial stereotactic-electroencephalography (SEEG) in patients suffering from drug-resistant epilepsy and implanted in both early visual and auditory areas for presurgical evaluation. SEEG reliably measures mesoscopic neural activity (Fukushima et al., 2015) with high spatial and temporal resolution, high resistance to possible artifacts (muscle contractions, eye blinks), and with an exceptional signal to noise ratio when compared to classic EEG methods used in humans (Lachaux et al., 2012).

In addition to the classical event-related potential analyses in the time domain, a useful theoretical framework to interpret neurophysiological activity at the mesoscopic level postulates that local changes in oscillations in different frequency bands encode different sensory/cognitive processes in a dynamic spectral fingerprint (Siegel et al., 2012). In line with this proposal, seminal studies have demonstrated associations between oscillations at different frequency bands and neurophysiological processes (Buzsáki and Wang, 2012, Engel and Fries, 2010, Kucewicz et al., 2014, Gregoriou et al., 2009). Of particular interest in the context of this research, SEEG allows the reliable recording of the neural activity in the high-gamma frequency band, an activity that has been linked to high-frequency synaptic and spiking activity (Manning et al., 2009, Buzsáki and Wang, 2012). In contrast, with non-invasive methods, such as magnetoencephalograhy (MEG) and electroencephalography (EEG), high-gamma power is often confounded by various sources of noise, such as ocular and muscle activity (Muthukumaraswamy, 2013, Carl et al., 2012).

In our study, we relied on the unique spatio-temporal resolution of SEEG to investigate how cross-modal processing (visual responses in auditory cortex or the reverse) and multisensory integration (alteration of the unimodal responses during bimodal stimulation) are reflected in intracranial event-related potentials (iERPs) and in power modulations of oscillatory activity during the first 150ms after stimulus onset.

## 2. Methods

### Participants

Because the objective of our study was to assess the presence of crossmodal inputs and multisensory integration in early/primary sensory cortices, we only included participants having electrodes implanted in both early temporal-auditory cortex (Heschl’s gyrus and planum temporale) and in early occipital-visual cortex (calcarine and pericalcarine regions). Even if implantations in both these sensory regions are rare, we managed to record from 47 electrodes in the temporal-auditory cortex and from 44 electrodes in the occipital-visual cortex in 8 patients.

The 8 participants (mean age: 34 years ± 11; 4 females) suffered from drug resistant epilepsy and were stereotactically implanted with intracerebral electrodes (Cardinale et al., 2012). Each electrode (diameter of 0.8 mm) comprised from 10 to 18 contacts (total number of contacts per patient: from 146 to 181, spaced 1.5 mm apart DIXI, Besançon, France). All patients had cognitive abilities in the normal range as assessed by a neuropsychological exam and did not have specific deficits in visual and auditory functions. Data were collected not before than three days after electrode implantation, 24 hours before and after spontaneous seizures, except for one patient experiencing one short seizure 2 hours before the acquisition and lasting less than 30 seconds, occurring with the recovery of the usual interictal intracranial EEG activity in less than 30 minutes. All the electrodes were implanted only according to clinical criteria, and the conduction of this study did not influence the clinical procedures. The research project was approved by the Institutional Review Boards of the Hospital Niguarda Ca’ Granda of Milan and adhered to the Declaration of Helsinki. The participants provided written informed consent.

### Procedure

The paradigm was implemented and administered using Presentation software (https://www.neurobs.com/). Each participant seated in front of the screen at a distance of 60 cm and was presented with several blocks (from 2 blocks to 12 blocks; median for all participants: 4 blocks) containing non-target and target auditory (A), visual (V) and audio-visual (AV) condition. A nothingcondition (N), without any stimulation, was used as control condition to record and compensate for any anticipatory brain responses at the times a stimulation is predicted to typically occur (Talsma and Woldorff, 2005, Gondan et al., 2005, Bonath et al., 2007, Teder-Sälejärvi et al., 2002). In every single block, each non-target and N conditions were presented 40 times, while each target condition was presented 4 times (10% occurrence). Non-target, target and N conditions were randomly interleaved. The V stimuli (subtending 14.8 deg. of visual angle) were presented for 100ms in the centre of the screen: the non-target V stimulus was a black and white checkerboard, while the target V stimulus was a coloured checkerboard. The A stimuli were presented binaurally through inserted earphones at a comfortable auditory level for each individual participant. The non-target A stimulus was a 100 ms segment of white noise (5 ms fade in/out), while the target a 100 ms pure tone (2000Hz; 5 ms fade in/out). The non-target AV condition was a combination of the non-target V and non-target A condition. During the non-target AV condition, the V stimulus was presented 30 ms before the onset of the A stimulus, since behavioural studies have shown that V stimuli presented before A stimuli (between 20 and 90 ms) allow obtaining the strongest behavioural gain during the audio-visual condition in comparison to the intramodal condition (Thorne et al., 2011); in agreement, neurophysiological studies have indicated that neural MSI in auditory areas occurs during a relatively extended temporal window and is particularly strong when the V stimulus precedes the A stimulus (between 30 and 75 ms) (Kayser et al., 2008, Musacchia and Schroeder, 2009, Thorne et al., 2011).

The inter-stimulus interval was jittered between 1000 and 1500 ms. The target AV condition was a combination of the target V and target A condition with the same temporal characteristic of the non-target AV stimulus. Timing accuracy of the presented stimuli was controlled offline with the Black Box toolkit (http://www.blackboxtoolkit.com). Participants were asked to maintain central fixation and to respond as fast as possible when the target conditions were presented. All patients were able to detect the target conditions with high accuracy (accuracy: 100% for all participants except for one (63%); false hits: from 0% to 7%, median 2%). We investigated the electrophysiological activity recorded during the non-target conditions in order to avoid any confounds in the signal that were linked to the motor response of the participant. Hereby, when mentioning A, V and AV conditions, we will refer to the respective input during the non-target conditions. Moreover, when mentioning the *intramodal* condition we will refer to the stimulus matching the sensory representations of the investigated cortex (i.e. auditory input in the auditory cortex, visual input in the visual cortex) (Stein and Stanford, 2008); while, when mentioning the *crossmodal* condition we will refer to the stimulus not matching the main sensory representations of the investigated cortex (i.e. auditory input in the visual cortex, visual input in the auditory cortex). Finally, when mentioning the *bimodal* condition we will refer to the combined intramodal and cross-modal inputs.

### Contacts of interest (COIs) localizations

We performed the analyses for contacts of interest (COIs) localized in Heschl’s gyrus and planum temporale (n=47) for the temporal-auditory cortex and in calcarine and pericalcarine regions (n=44) for the occipital-visual cortex. To identify COIs in each participant, post-implantation intraoperative Cone-Beam-CT (CBCT) scan (192 axial slices, 512 x 512 matrix, 0.415 x 0.415 x 0.833 mm anisotropic voxels) was registered to preimplantation MR (3D fast field echo T1-weighted sequence, contiguous axial slices with 560 x 560 reconstruction matrix, 0.46 x 0.46 x 0.9 mm voxel, no inter-slice gap) by means of FLIRT 6.0 (Jenkinson and Smith, 2001). Such CBCT scans provide undistorted images of the electrodes (figure 1). The anatomical location of COIs was assessed with a two-steps procedure: 1) an expert medical epileptologist (R.M.) performed a visual inspection of the co-registered images in the native MRI space with 3D Slicer 4.3.1 software (Fedorov et al., 2012), and identified possible COIs; 2) by means of Freesurfer 5.3.0 (Fischl, 2012), normalized brain tissue was segmented and cerebral surfaces were reconstructed and parcellated. Electrodes located in the visual (i.e. calcarine and pericalcarine) and auditory regions of interest (i.e. Heschl’s gyrus and planum temporale) were identified as COIs.

**Figure 1:**
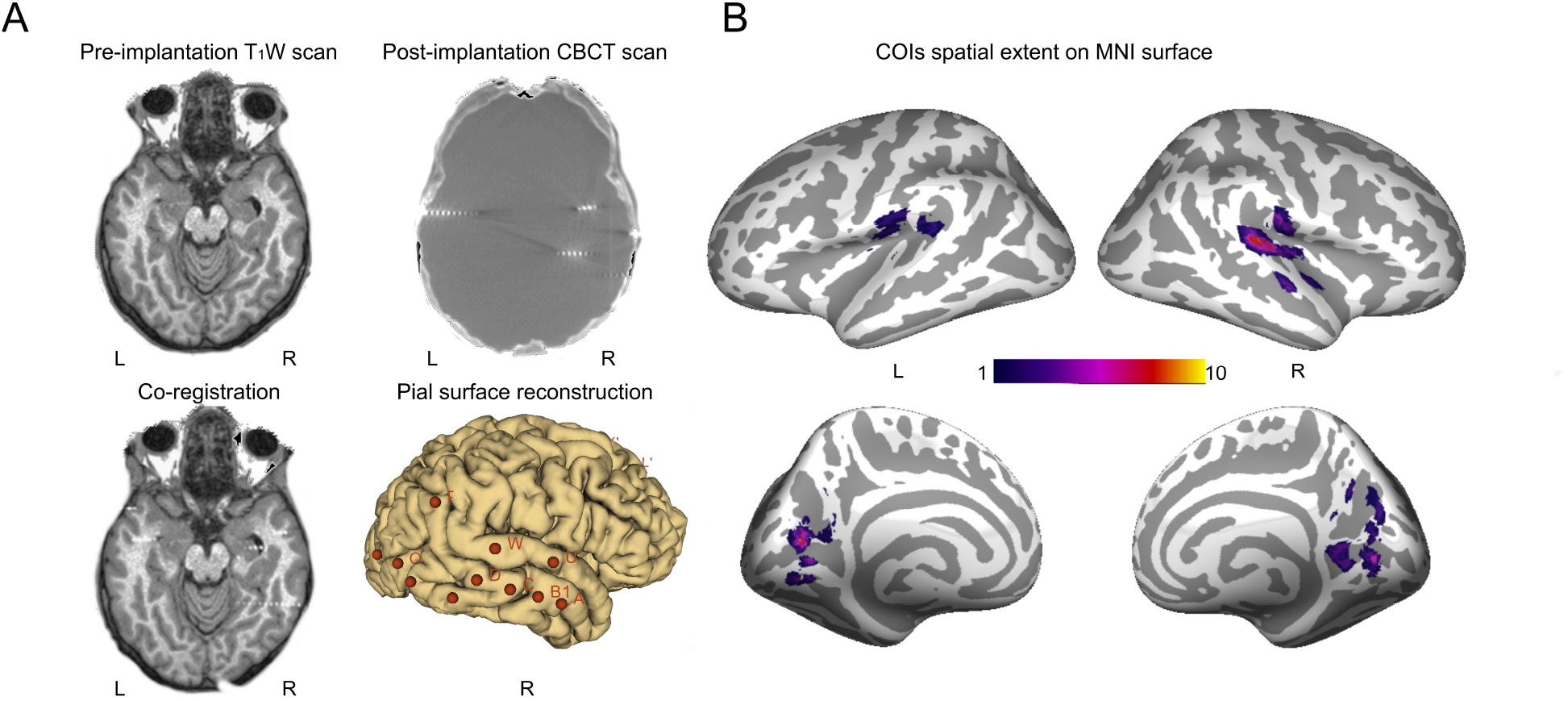
(A) Exemplar of pre-implantation T1W scan, post-implantation CBCT scan and simultaneous visualization of both co-registered datasets from one of the studied subjects. From the same participant, pial surface reconstruction with fiducial markups at electrode entry points is also depicted; (B) Spatial extension of COIs from all participants superimposed on MNI surface (auditory cortex: n = 47; visual cortex n 44). COIs = contacts of interest; L = left; R = right.

### SEEG recordings and signal pre-processing

Continuous SEEG was recorded by means of EEG-1200 Neurofax (Nihon Kohden), comprising 192 channels (1000 Hz sampling rate). A medical epileptologist (R.M.) visually inspected the raw signal and did not detect any pathological activity in any investigated COI. We performed all the analyses with Fieldtrip software package (http://www.fieldtriptoolbox.org/) using the bipolar montage; therefore, our results are referred to 36 couples of COIs in the auditory cortex and 35 in the visual cortex. Bipolar montage represents a conservative approach since a source which activity is similarly recorded by two contiguous contacts will be removed by the application of the bipolar montage (Zaveri et al., 2006). However, we chose to use this montage because volume conduction phenomena, heavily affecting non-invasive methods (e.g. magnetoencephalograhy and electroencephalography), can also possibly affect SEEG (Kajikawa and Schroeder, 2011), making it difficult to assert that the integrative process truly occurs in early sensory region since proximate associative regions also engage in MSI. Therefore, the use of a bipolar montage allows us to ascertain that potential crossmodal/multisensory effects truly originate from our selected early sensory regions where the COIs are located.

The raw signal was down-sampled at 500 Hz. Each trial was detrended and epoched in a time-window of 900 ms preto 900 ms post-stimulus onset. Trials showing artifacts were removed. We analysed the SEEG signal in the time-frequency domain investigating the power in the different frequency bands. Given that the timing of the neurophysiological emergence of cross-modal or MSI modulation is important to interpret early MSI mechanisms and that the temporal resolution in the timefrequency domain presents a certain degree of uncertainty (Cohen, 2014), we performed the analyses also in the time-domain, investigating intracranial event-related potentials (iERPs).

### iERPs analyses

In each single subject, the pre-processed signal was baseline corrected (0.200-0.050s pre-stimulus onset), low-pass filtered (30 Hz) and averaged across trials for each condition of interest (A, V, AV and N conditions). Between-conditions differences were assessed comparing the post-stimulus amplitude (0-150ms post-stimulus onset) between the relevant conditions. To test for significance, the conditions of interest were compared using two-tailed independent samples t-tests (p¡0.05 FDR-corrected). We assessed the presence of: 1) *intramodal* processing, comparing the intramodal input with the control ‘nothing’ condition (A vs. N in the auditory cortex; V vs. N in the visual cortex); 2) *cross-modal* processing, comparing the cross-modal input with the control ‘nothing’ condition (V vs. N in the auditory cortex; A vs. N in the visual cortex); 3) *multisensory* processing, comparing the responses to the bimodal input with the intramodal input eliciting the maximum response in that cortex (AV vs. A in the auditory cortex; AV vs. V in the visual cortex), in agreement with the maximum model (Stein and Meredith, 1993). Any COI showing intramodal processing, was classified as functional COI (fCOI). fCOIs were labelled as: 1) *unimodal fCOIs*, when they presented only intramodal processing; 2) *bimodal fCOIs* when they presented both intramodal and crossmodal processing but no multisensory processing; 3) *MSI fCOIs* when they presented multisensory processing. When bimodal fCOIs and MSI fCOIs were identified, we verified if the observed multisensory activity was additive or non-additive (Giard and Peronnet, 1999, Molholm et al., 2002), comparing the linear summation of the responses to V and A (V+A) conditions with the linear summation of the responses to AV and N (AV+N) conditions, by means of a two-tailed paired-samples t-test (p < 0.05 FDR-corrected). The N condition was used to control for any anticipatory brain responses or unknown cognitive factors that would be summed twice in both the factors of the equation (V+A vs. AV+N) (Teder-Sälejärvi et al., 2002, Talsma and Woldorff, 2005).

### Time-frequency analyses: power domain

In each subject, time-frequency analyses were performed by convolving the pre-processed signal of each individual trial with complex Morlet wavelets in the time-window from 900 ms pre to 900 ms post-stimulus onset in steps of 10 ms. Signal decomposition was performed in two different frequency-windows: 4-30 Hz and 30-200 Hz in order to optimize the trade-off between temporal and frequency precision (Cohen, 2014). For the 4-30 Hz frequency window, the power values of each condition of interest (i.e. A, V, AV, and N) were estimated with wavelet widths ranging from 4 to 5 cycles in steps of 1 Hz. For the 30-200 Hz frequency window, the power values were estimated with wavelet widths ranging from 5 to 10 cycles in steps of 5 Hz. For both frequency-windows, the wavelet widths changed linearly as a function of frequency. The obtained power values were then normalized by means of a logarithmic transformation in order to apply parametric statistics (Kiebel et al., 2005). Statistical analyses were then performed in a temporal window of 150ms after stimulus onset and separately for the *θ* (4-8 Hz), *α* (8-13 Hz), *β* (13-30 Hz), *γ* (30-80 Hz), and high-*γ* band (80-200 Hz). The conditions of interest were compared using two-tailed independent samples t-tests. In each COI and for each frequency band, the results were FDRcorrected (p¡0.05) for the observed time-frequency points. As for the iERPs, we assessed the presence of intramodal, crossmodal and multisensory processing. Any COI showing intramodal processing across any frequency band was classified as functional COI (fCOI). Then, in each single frequency band, fCOIs were labelled as: 1) *unimodal fCOIs*, when they presented only intramodal processing; 2) *bimodal fCOIs* when they presented both intramodal and cross-modal processing but no multisensory processing; 3) *cross-modal fCOIs* when they presented only cross-modal processing; 4) *MSI fCOIs* when they presented multisensory processing. When MSI fCOIs were identified, we verified whether the observed multisensory activity was additive or non-additive. Due to the non-linear properties of the wavelet transformed oscillatory responses, we used the procedure proposed by Senkwoski et al. (2007) (Senkowski et al., 2007). Each single pre-processed epoch of the V condition was summed with all the preprocessed epochs of the A condition to obtain the V+A epochs; the same was implemented for AV with the N condition (AV+N epochs). Each obtained V+A epoch and AV+N epoch was then convolved with complex Morlet wavelets using the same parameters used for the time-frequency analyses described above. Subsequently, in each wavelet-transformed epoch and separately for V+N and AV+N condition, we extracted the maximum peak of the activity in the time-frequency window in which we have observed the significant multisensory activity. A bootstrap procedure was then implemented: peaks were randomly selected separately for V+A epochs and AV+N epochs. The number of these randomly selected peaks was equal to the number of the original trial of the AV condition. This procedure was repeated 10000 for each V+A and AV+N condition. For each of these bootstrapped samples, we computed the mean of the peaks. Then, for the AV+N condition, we computed the mean of the sample means of the peaks. This mean was compared with the distribution of the sample means of the peaks obtained from the bootstrap procedure of V+A condition. The percentile of the AV+N condition relative to the bootstrap sample means was computed. If it was below 2.5% or above 97.5%, AV+N was considered significantly different from the A+V condition and therefore multisensory integration was considered respectively, subadditive or superadditive (Stein et al., 2009).

## 3. Results

We considered evidence of MSI the presence of fCOIs classified as *MSI fCOIs* (auditory cortex: AV vs. A; visual cortex: AV vs. V), *bimodal fCOIs* (auditory cortex: V vs. N; visual cortex: A vs. N; with evidence of intramodal processing in the same frequency band for the power analyses) and *cross-modal fCOIs* (auditory cortex: V vs. N; visual cortex: A vs. N; with evidence of intramodal processing across the whole spectrum, but not in the same frequency band for the time-frequency analyses).

### iERPs results

A summary of the following results is presented in figure 2.

**Figure 2:**
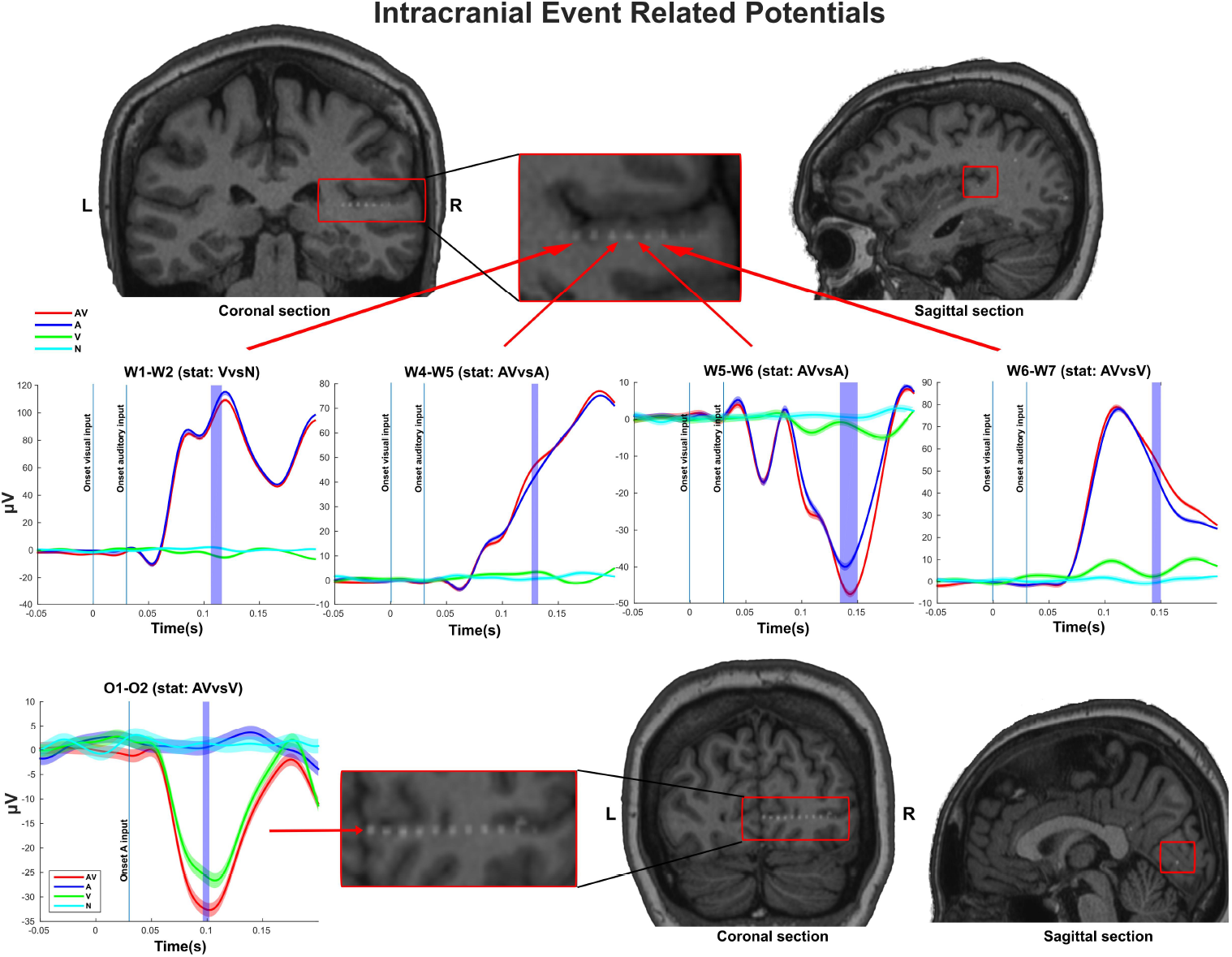
(A) Intracranial event-related potentials (iERPs) of functional contacts of interest (fCOIs) during auditory (in blue), visual (in green), audiovisual (in red) and ‘nothing’ stimulation. In purple the significant time points showing evidence of multisensory integration as assessed by the relevant statistic (AV vs. A and V vs. N in the auditory cortex; AV vs. V in the visual cortex).MRI images with superimposed CBCT scan show the localization of the relevant fCOIs (top figure: Heschl’s gyrus and planum temporale; bottom figure: calcarine scissure).

#### Auditory cortex

In the auditory cortex, 86% of COIs (31 out of 36 COIs) presented significant responses to the auditory stimulation and, therefore, they were classified as fCOIs. MSI effects, detected as bimodal or MSI fCOIs (see Methods section), were observed in 4 fCOIs (13%) of the auditory cortex. All these fCOIs belonged to the same intracranial electrode of a single participant: one of these fCOIs was located in right Heschl’s gyrus, while the remaining ones were contiguous and were located in the right planum temporale. In these fCOIs, the first effect of the intramodal processing occurred in Heschl’s gyrus at around 16 ms after the auditory input onset, while the first MSI effect, occurred around 106ms after the onset of the visual stimulation and it was detected in a bimodal fCOI. Then, MSI appeared in the three more centrifugal fCOIs, labelled as MSI fCOIs, at 126ms, 134ms and 142ms after the visual onset of the AV condition. These fCOIs presented enhanced or depressed responses (figure 3), but with no evidence of non-additive MSI.

**Figure 3:**
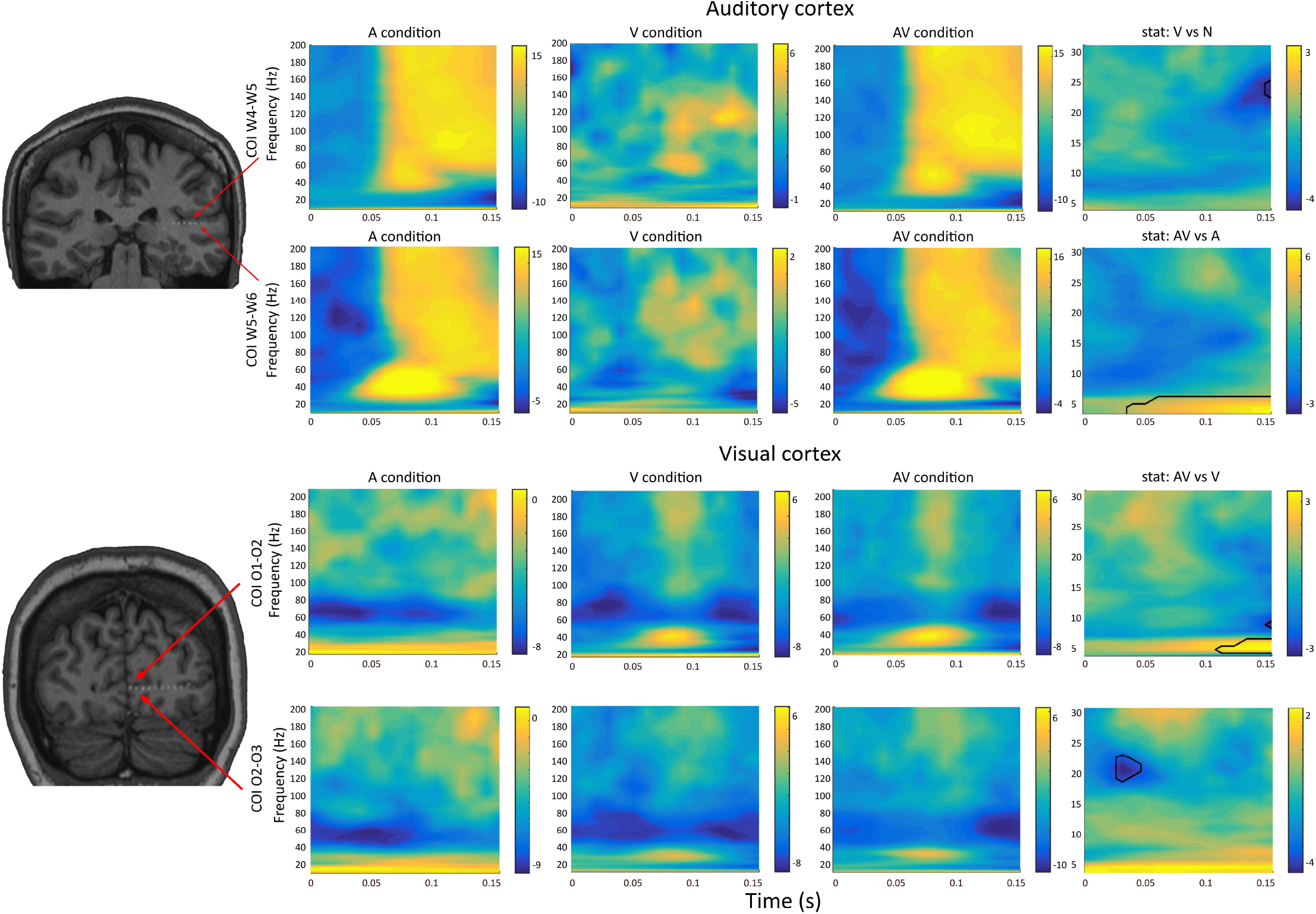
Time-frequency analyses: response exemplars of functional contacts of interest (fCOIs) during the auditory, visual and audio-visual condition expressed as baseline corrected z-scores in the relative fCOI. In the last column t-scores representations of the relevant statistics with solid black lines representing significant responses. MRI images with superimposed CBCT scan show the localization of the fCOIs (top figure: Heschl’s gyrus; bottom figure: calcarine scissure).

#### Visual cortex

In the visual cortex, 29% of COIs (10 out of 35 COIs) presented significant responses to the visual stimulation and, therefore, they were classified as fCOIs. MSI was observed in 1 fCOIs (10%). The earliest intramodal processing effect was detected at around 74 ms after the visual input processing, while the earliest MSI effect was detected in one MSI fCOI, at 68ms after the auditory input onset of the AV condition. This MSI effect was evident as a depression of the iERPs (figure 3), with no evidence of non-additive MSI.

### Time-frequency domain: power results

A summary of these results is presented in figures 3 and 4 and table 1.

**Table 1:** Time-frequency domain power results: (*) percentages expressed over fCOIs exhibiting intramodal activity in that frequency band; (^) percentages expressed over all fCOIs across the whole investigated spectrum. (Abbreviations: AUD = auditory cortex, VIS = visual cortex, fCOI = functional contact of interest).

| | 4-8Hz($\theta$) | | 8-13Hz($\alpha$) | | 13-30Hz($\beta$) | | 30-80Hz($\gamma$) | | 80-200Hz(high-γ) | |
| --- | --- | --- | --- | --- | --- | --- | --- | --- | --- | --- |
|  | Aud | Vis | Aud | Vis | Aud | Vis | Aud | Vis | Aud | Vis |
| *fCOIs\** | 29 (81%) | 8 (23%) | 31 (86%) | 12 (34%) | 33 (92%) | 11 (31%) | 36 (100%) | 11 (31%) | 35 (97%) | 11 (31%) |
| *Unim fCOIs\*\** | 27 (93%) | 7 (88%) | 31 (100%) | 8 (67%) | 32 (97%) | 9 (82%) | 36 (100%) | 10 (91%) | 35 (100%) | 11 (100%) |
| *Bim fCOIs\*\** | 1 (3%) | 0 (0%) | 0 (0%) | 2 (17%) | 1 (3%) | 0 (0%) | 0 (0%) | 0 (0%) | 0 (0%) | 0 (0%) |
| *MSI fCOIs\*\** | 1 (3%) | 1 (13%) | 0 (0%) | 2 (17%) | 0 (0%) | 2 (18%) | 0 (0%) | 1 (9%) | 0 (0%) | 0 (0%) |
| *Bim fCOIs^* | 1 (3%) | 0 (0%) | 0 (0%) | 2 (13%) | 1 (3%) | 0 (0%) | 0 (0%) | 0 (0%) | 0 (0%) | 0 (0%) |
| *MSI fCOIs^* | 1 (3%) | 1 (6%) | 0 (0%) | 2 (13%) | 0 (0%) | 2 (13%) | 0 (0%) | 1 (6%) | 0 (0%) | 1 (6%) |
| *Crossmodal fCOIs^* | 0 (0%) | 0 (0%) | 0 (0%) | 0 (0%) | 0 (0%) | 0 (0%) | 0 (0%) | 0 (0%) | 0 (0%) | 3 (19%) |

**Figure 4:**
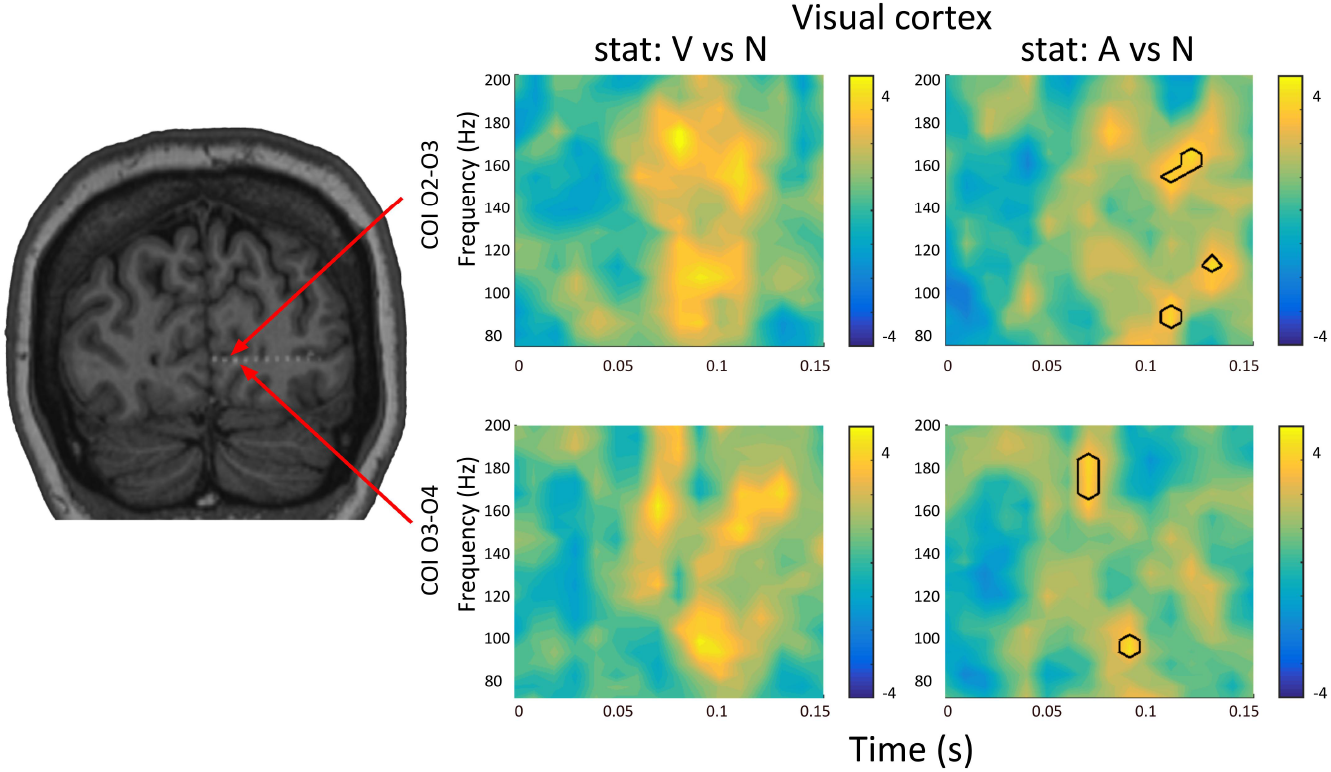
Time-frequency analyses: response exemplars of functional contacts of interest (fCOIs) located in the early visual cortex showing evidence of significant high-*γ* power modulations during the cross-modal input and with no significant modulations during the intramodal stimulation in the same frequency band. Power modulations are represented as t-scores of the relevant statistics. Solid black lines represent significant responses.

#### Auditory cortex

In the auditory cortex, all COIs (36 out of 36) presented intramodal processing in the whole investigated spectrum and therefore they were classified as fCOIs (total fCOIs). There was evidence of intramodal processing also in the majority of the COIs (from 81% to 100%) when considering each single frequency band (*θ, α, β, γ* and high-*γ*). We observed that 2 fCOIs (6% of the total fCOIs) presented cross-modal processing: 1 (3%) in the *θ* band, with an enhanced activity, and 1 (3%) in the *β* band, with a depressed activity. In the same frequency bands, these fCOIs presented also intramodal processing, therefore, they were labelled as bimodal fCOIs. Multisensory processing was observed in the *θ* band in 1 fCOIs (3% of total fCOIs), as a superadditive activity. In the same frequency band, this fCOI also presented intramodal processing, therefore it was labelled as MSI fCOIs. The observed bimodal and MSI fCOIs were located in the right planum temporale of the same participant and they were contiguous: the bimodal fCOIs were more centripetal in respect to the MSI fCOI.

#### Visual cortex

In the visual cortex, 46% of COIs (16 out of 35) showed intramodal processing across the whole spectrum and, therefore, they were classified as fCOIs. Intramodal processing was also observed in each single frequency band, although to a lesser degree in comparison to the auditory cortex (from 23% of fCOIs in the *θ* band to 34% of COIs in the *α* band).

We observed that 1 fCOI (6% of the total fCOIs) presented cross-modal processing in the *α* band, with a depressed activity: this fCOI also showed intramodal processing in the same frequency band and, therefore, it was labelled as bimodal fCOIs. Remarkably, cross-modal processing was also observed in 3 fCOIs (19% of total fCOIs) in the high-*γ* band with enhanced responses; however, in these fCOIs the intramodal processing occurred in the low-frequency bands, but not in the high-*γ* band. These fCOIs were, therefore, classified as *cross-modal fCOIs*.

*Multisensory processing* was detected in 1 fCOIs (6% of total fCOIs) in the *θ* band (showing enhanced and additive activity), in 2 fCOIs (13% of total fCOIs) in the *α* band (showing enhanced and subadditive activity), in 2 fCOIs (13% of total fCOIs) in the *β* band (showing depression with both additive and subadditive activity) and in 1 fCOIs (6% of total fCOIs) in the *γ* band (showing enhanced but additive activity). Due to the presence of both intramodal and multisensory processing in the same frequency band, all these fCOIs were labelled as MSI fCOIs. Multisensory processing was also observed in the high-*γ* band in 1 fCOI (6% of total fCOIs): in this case, as observed for the cross-modal fCOIs, the multisensory processing was not accompanied by the intramodal processing in the same frequency band.

## 4. Discussion

In our study, we capitalized on the unique spatio-temporal resolution of SEEG recordings to investigate the involvement of the human early visual (calcarine and pericalcarine regions) and auditory (Heschl’s gyrus and planum temporale) cortices in the processing of cross-modal inputs and multisensory integration. To this end, we investigated the SEEG signal in the time domain (investigating iERPs) and time-frequency domain (investigating power), during an audio-visual paradigm in the first 150ms post-stimulus onset.

The iERPs results evidenced MSI effects that occurred earlier in the visual cortex (starting at 68ms after the auditory onset) than in the auditory cortex (starting from 106ms after the visual onset). This temporal effect is congruent with the fact that auditory input is processed earlier in the auditory cortex (starting at 16ms in our study) compared to the visual input in the visual cortex (starting at 74ms in our study). Our results are also in agreement with the previous literature showing that MSI may occur at 80-90ms in the human auditory cortex and at 40-50ms in the visual cortex during AV integration (Giard and Peronnet, 1999, Molholm et al., 2002, Raij et al., 2010); while the earliest intramodal processing effects were described to occur within the first 15ms (Molholm et al., 2002) in the auditory cortex and around 40-60ms in the visual cortex (Foxe et al., 2002, Raij et al., 2010). Such temporal profiles suggest that each sensory input is first processed in its dominant sensory cortex and then conveyed to the ectopic sensory region (Raij et al., 2010). The transfer of information between sensory cortices might rely on existing monosynaptic projections connecting in both directions the early auditory and visual cortices (Falchier et al., 2002, Rockland and Ojima, 2003, Hall and Lomber, 2008, Cappe and Barone, 2005, Bizley et al., 2006).

Our time-domain results showed another interesting feature: in the early auditory cortex, MSI effects are present with a temporal gradient from the most centripetal fCOI (bimodal processing at 106ms), in the Heschl’s gyrus, to the most centrifugal fCOIs in the planum temporale (MSI processing at 126ms, 134ms and 142ms). This phenomenon suggests that the primary source of the MSI effects is located in the Heschl’s gyrus, potentially supporting the observation that the primary auditory cortex is the target of projections from the visual cortex (Budinger et al., 2006, Cappe and Barone, 2005), and in agreement with the notion that the core regions of the auditory cortex present the shortest latencies during auditory tasks in comparison to the growing latencies of the other auditory regions (Recanzone et al., 2000).

The time-frequency results extended the iERPs findings by identifying markers of cross-modal processing and MSI in the local power changes across different spectral bands. In the early auditory cortex, MSI effects were evident in the low-frequency bands (*θ* and *β* bands), while, in the early visual cortex, MSI effects were evident across all the frequency bands, with a dominant expression in the *α* and in the high-*γ* band. We found evidence of both additive and subadditive MSI activity in the visual cortex, while only superadditive MSI effects were observed in the auditory cortex.

Although we still lack of a comprehensive understanding of the link between oscillatory activity in different frequency bands and specific neurophysiological mechanisms, several findings have provided evidence that different frequencies of coherent oscillations might preside over the directions of information flow, suggesting that the brain segregates information in different frequency channels (Wang, 2010, Buffalo et al., 2011). For instance, the superficial layers of the visual cortex in monkeys present a strong synchronization in the *γ* band, while the deep layers rely more on the *α*/*β* band (Buffalo et al., 2011). The authors argued that the synchronization in these two frequency bands is transferred to different targets based on the long-standing notion that the superficial layers (mainly layers 2/3) implement feedforward projections, while the deep-layers implement feedback projections. Moreover, it was shown that feedforward modulations are carried by the theta and the *γ* band while feedback influences by the *β* band in the visual cortex (Bastos et al., 2015). Similarly, simultaneous recording from V1 and V4 confirmed that the *γ* oscillations propagated in feedforward direction, while the *α* oscillations in the feedback direction (Van Kerkoerle et al., 2014). Taken together, these results suggest precise inter-areal information flow by means of specific frequency channels: the low frequency bands (in particular the *α* and *θ* rhythms) seem to preside over the feedback interactions, while the *γ* band over feedforward interactions (Fries, 2015, Zheng and Colgin, 2015). Based on the previous evidence of the presence of direct (monosynaptic) heteromodal connections between the early auditory and visual areas (Cappe et al., 2009, Falchier et al., 2002, Hall and Lomber, 2008, Rockland and Ojima, 2003), it is plausible that these mechanisms of intramodal interactions are at work also for cross-modal interactions. In our study, both the investigated sensory cortices presented power modulations mainly in the low-frequency bands (*θ* and *β* bands in the auditory cortex and mainly the *α* band for the visual cortex), suggesting that feedback cortico-cortical mechanisms might govern MSI in early sensory cortices (Bastos et al., 2015, Fontolan et al., 2014, Van Kerkoerle et al., 2014).

Previous studies, using intracranial electrophysiological recordings in humans, showed that crossmodal inputs can induce pure phase reset of the low frequency oscillatory activity in the visual (Mercier et al., 2013) and auditory (Mercier et al., 2015) cortices. Such pure phase resetting suggests that ectopic inputs can induce a reset of the phase of the intracranial signal without modulation of the power activity (Lakatos et al., 2007). Differently from these studies, we found that crossmodal inputs induce low-frequency power modulations in both sensory cortices (*θ* band in the auditory cortex and *α* band in the visual cortex). Such inconsistencies should be carefully investigated in future studies in the light of the possible different statistical properties of the phase and power of the oscillatory activity in the time-frequency domain (Ding and Simon, 2013).

Another important result of our study is the presence in the early visual cortex of power modulation in the high-*γ* band by sounds (see fig.4). High-*γ* activity is considered a marker of synaptic and spiking activity (Manning et al., 2009, Buzsáki and Wang, 2012). Remarkably, all these visual fCOIs showing increase in high-*γ* activity by sounds did not present high-*γ* band power modulations during intramodal/visual processing, which was evident only in the low-frequency bands. These observations suggest: 1) the presence of auditory responsive neuronal populations in early visual areas and that 2) the visual and auditory responses of these neuronal populations are not in spatial register with each other since our sounds and visual stimuli were originating from the same position in external space. This possibility is supported by the notion that monosynaptic connections from the early auditory cortex project mainly to the peripheral visual fields (Falchier et al., 2002, Hall and Lomber, 2008) where our visual-only stimuli, presented in foveal position, did not elicit any apparent neuronal activity. Alternatively, this observation might suggest the presence of auditory only responsive neuronal populations in human early visual areas, but we find this explanation unlikely. One of the rare previous human study relying on intracranial recording did not find power modulation in the high-*γ* band in the primary visual cortex during MSI (Quinn et al., 2014). Such inconsistency with our observation of high-*γ* modulation by sounds in the occipital cortex could relate to the fact that we investigated audiovisual integration while Quinn and collaborators (2013)(Quinn et al., 2014) investigated visual-tactile integration, suggesting that audio-visual or visuo-tactile integration may rely on different neurophysiological mechanisms in the primary visual cortex.

**In summary**, by analysing the electrophysiological signal both in the time and frequency domain, our results support the idea that MSI occurs at the earliest stages of the sensory processing hierarchy (Foxe et al., 2002, Ghazanfar and Schroeder, 2006), potentially through direct anatomical connections between the visual and auditory cortices. Moreover, our study compellingly illustrate how stereotactic electrophysiological recording in humans represents a unique technique to investigate the multisensory nature of brain regions, in particular those classically considered unisensory.

## Acknowledgment

The Authors are grateful to Prof. Thomas Thesen for his insightful comments during the data analyses.

